# Immunization with the receptor–binding domain of SARS-CoV-2 elicits antibodies cross-neutralizing SARS-CoV-2 and SARS-CoV without antibody-dependent enhancement

**DOI:** 10.1101/2020.05.21.107565

**Authors:** Jinkai Zang, Chenjian Gu, Bingjie Zhou, Chao Zhang, Yong Yang, Shiqi Xu, Xueyang Zhang, Yu Zhou, Lulu Bai, Yang Wu, Zhiping Sun, Rong Zhang, Qiang Deng, Zhenghong Yuan, Hong Tang, Di Qu, Dimitri Lavillette, Youhua Xie, Zhong Huang

**Author notes:** These authors contributed equally (Jinkai Zang, Chenjian Gu, Bingjie Zhou, Chao Zhang, Yong Yang). Corresponding author: Zhong Huang or Youhua Xie or Dimitri Lavillette.

## Abstract

Recently emerged severe acute respiratory syndrome coronavirus 2 (SARS-CoV-2) is the pathogen responsible for the ongoing coronavirus disease 2019 (COVID-19) pandemic. Currently, there is no vaccine available for preventing SARS-CoV-2 infection. Like closely related severe acute respiratory syndrome coronavirus (SARS-CoV), SARS-CoV-2 also uses its receptor-binding domain (RBD) on the spike (S) protein to engage the host receptor, human angiotensin-converting enzyme 2 (ACE2), facilitating subsequent viral entry. Here we report the immunogenicity and vaccine potential of SARS-CoV-2 RBD (SARS2-RBD)-based recombinant proteins. Immunization with SARS2-RBD recombinant proteins potently induced a multi-functional antibody response in mice. The resulting antisera could efficiently block the interaction between SARS2-RBD and ACE2, inhibit S-mediated cell-cell fusion, and neutralize both SARS-CoV-2 pseudovirus entry and authentic SARS-CoV-2 infection. In addition, the anti-RBD sera also exhibited cross binding, ACE2-blockade, and neutralization effects towards SARS-CoV. More importantly, we found that the anti-RBD sera did not promote antibody-dependent enhancement of either SARS-CoV-2 pseudovirus entry or authentic virus infection of Fc receptor-bearing cells. These findings provide a solid foundation for developing RBD-based subunit vaccines for SARS-CoV2.

## INTRODUCTION

In December 2019, the coronavirus disease 2019 (COVID19), caused by a novel coronavirus named severe acute respiratory syndrome coronavirus 2 (SARS-CoV-2), was first identified in Wuhan, China (Zhou et al., 2020; Zhu et al., 2020). Since then, COVID19 cases have been increasingly reported in many countries and on March 11, 2020, the World Health Organization (WHO) declared COVID19 as a pandemic. Although the symptoms associated with COVID19 are generally mild, approximately 20% of COVID19 patients may develop severe clinical manifestations such as pneumonia, acute respiratory distress syndrome, sepsis and even death (Wu and McGoogan, 2020). According to WHO, as of April 30, 2020, there are 3,090,445 confirmed COVID19 cases and 217,769 deaths in total in the world (https://www.who.int/docs/default-source/coronaviruse/situation-reports/20200430-sitrep-101-covid-19.pdf?sfvrsn=2ba4e093_2). Clearly, COVID19 is a serious public health crisis. However, there is currently no vaccine available for COVID19.

SARS-CoV-2 belongs to the genus *Betacoronavirus* of the family *Coronaviridae* (Gorbalenya et al., 2020). Like other coronaviruses, SARS-CoV-2 is an enveloped virus and possesses a ~30 kb single-stranded positive-sense RNA genome. This viral genome encodes 4 structural proteins including spike (S), envelope (E), membrane (M), and nucleocapsid (N) proteins, 16 nonstructural proteins, and a few accessory proteins (Wu et al., 2020). The S protein consists of an ectodomain, a transmembrane domain, and a short intracellular tail (Walls et al., 2020). The ectodomain can be further divided into two functionally distinct subunits, S1 and S2, which are responsible for receptor binding and membrane fusion, respectively. Like the closely related severe acute respiratory syndrome coronavirus (SARS-CoV), SARS-CoV-2 also uses human angiotensin-converting enzyme 2 (ACE2) as the key receptor to facilitate its entry into host cells (Zhou et al., 2020). The S protein binds human ACE2 protein through its receptor-binding domain (RBD) located within the S1 subunit (Lan et al., 2020; Shang et al., 2020; Wang et al., 2020b; Yan et al., 2020)

Many approaches have been tested for rapid development of SARS-CoV-2 vaccines, yielding some exciting results (Amanat and Krammer, 2020; Chen et al., 2020). For example, Gao et al. recently reported that an inactivated whole-virus vaccine provided protection in macaques against experimental SARS-CoV-2 challenge (Gao et al., 2020). Thus far, a number of SARS-CoV-2 vaccine candidates derived from different vaccine platforms, including DNA vaccine, mRNA vaccine, inactivated whole virus vaccine, and adenovirus-vectored vaccine, have rapidly progressed into clinical trials (Amanat and Krammer, 2020; Callaway, 2020; Chen et al., 2020).

One of the challenges in developing vaccines for coronaviruses is a potential vaccine-induced immune enhancement of disease (Hotez et al., 2020; Huisman et al., 2009). For example, immunization with inactivated whole-virus SARS-CoV vaccine was found to elicit an immune response that exaggerate disease upon viral challenges in animal models (Bolles et al., 2011; Luo et al., 2018; Tseng et al., 2012; Wang et al., 2016). Results from some other studies suggest that antibodies targeting the spike protein of coronaviruses play a major role in antibody-dependent enhancement (ADE) likely through increasing the binding/entry of antibody-bound virion to Fc receptor (FcR)-expressing cells (Corapi et al., 1992; Jaume et al., 2011; Liu et al., 2019; Olsen et al., 1992; Wan et al., 2020; Wang et al., 2014; Yip et al., 2014). So far, all the SARS-CoV-2 vaccine candidates entering clinical trials contain or express full-length or near full-length S protein and therefore bear the risk of ADE. Thus, it is important to continue the search for a safe and effective SARS-CoV-2 vaccine.

Recombinantly produced RBD proteins of SARS-CoV and MERS-CoV have been shown to potently induce protective neutralizing antibodies against respective viruses in preclinical studies and are therefore considered promising vaccine candidates (reviewed in (Jiang et al., 2012; Zhou et al., 2018)). In the present study, we investigated the vaccine potential of SARS-CoV-2 RBD (hereafter referred as SARS2-RBD). We found that immunization of mice with recombinant SARS2-RBD elicited the production of serum antibodies that efficiently neutralized both SARS-CoV-2 and SARS-CoV but did not promote ADE.

## RESULTS

### Mice immunized with RBD-Fc fusion protein produced SARS-CoV-2 neutralizing antibodies

To rapidly evaluate the vaccine potential of SARS2-RBD, a pilot mouse immunization study was performed with a recombinant RBD fusion protein containing the Fc region of mouse IgG1 at the C-terminus (RBD-Fc) as the immunogen. The mice received three vaccine doses at days 0, 8, and 13, respectively. The first vaccine dose contained 100 μg of RBD-Fc protein, 500 μg of aluminum hydroxide and 25 μg of CpG, the second one contained 50 μg of RBD-Fc in complete Freund’s adjuvant, and the third one contained 50 μg of RBD-Fc formulated with Titermax adjuvant. One week after the last immunization (day 20), serum samples were collected from the three immunized mice for antibody measurement. All three antisera dose-dependently reacted with His-tagged SARS2-RBD in ELISA, whereas the control sera from an age-matching naïve mouse did not show significant reactivity regardless of the sera doses used (Fig. 1A). The anti-RBD-Fc sera #1 that had the highest RBD-binding titer (2×10^5^) was selected for further functional analyses. The anti-RBD-Fc sera #1 could dose-dependently inhibit the binding between recombinant ACE2-Fc fusion protein and His-tagged SARS2-RBD in competition ELISA (Fig. 1B), indicating that the antisera contain antibodies targeting the ACE2-binding motif within RBD. The anti-RBD-Fc sera #1 was then assessed for its ability to neutralize SARS-CoV-2 S protein-pseudotyped retrovirus (hereafter referred as SARS2-PV) and to neutralize live SARS-CoV-2. As shown in Fig. 1C, the antisera dose-dependently neutralized SARS2-PV entry with a calculated NT50 value of 10513. Moreover, addition of the anti-RBD-Fc sera #1 prevented the development of cytopathic effect (CPE) in SARS-CoV-2-inoculated VeroE6 cells (Supplementary Fig. 1). Results from qRT-PCR and IFA assays showed that the anti-RBD-Fc sera #1 were highly effective, neutralizing >50% infection even at the serum dilution of 1:5120 (Fig. 1D-E). Taken together, the above results demonstrate that RBD-Fc is an immunogen capable of efficiently inducing SARS-CoV-2-neutralizing antibodies.

**Figure 1.**
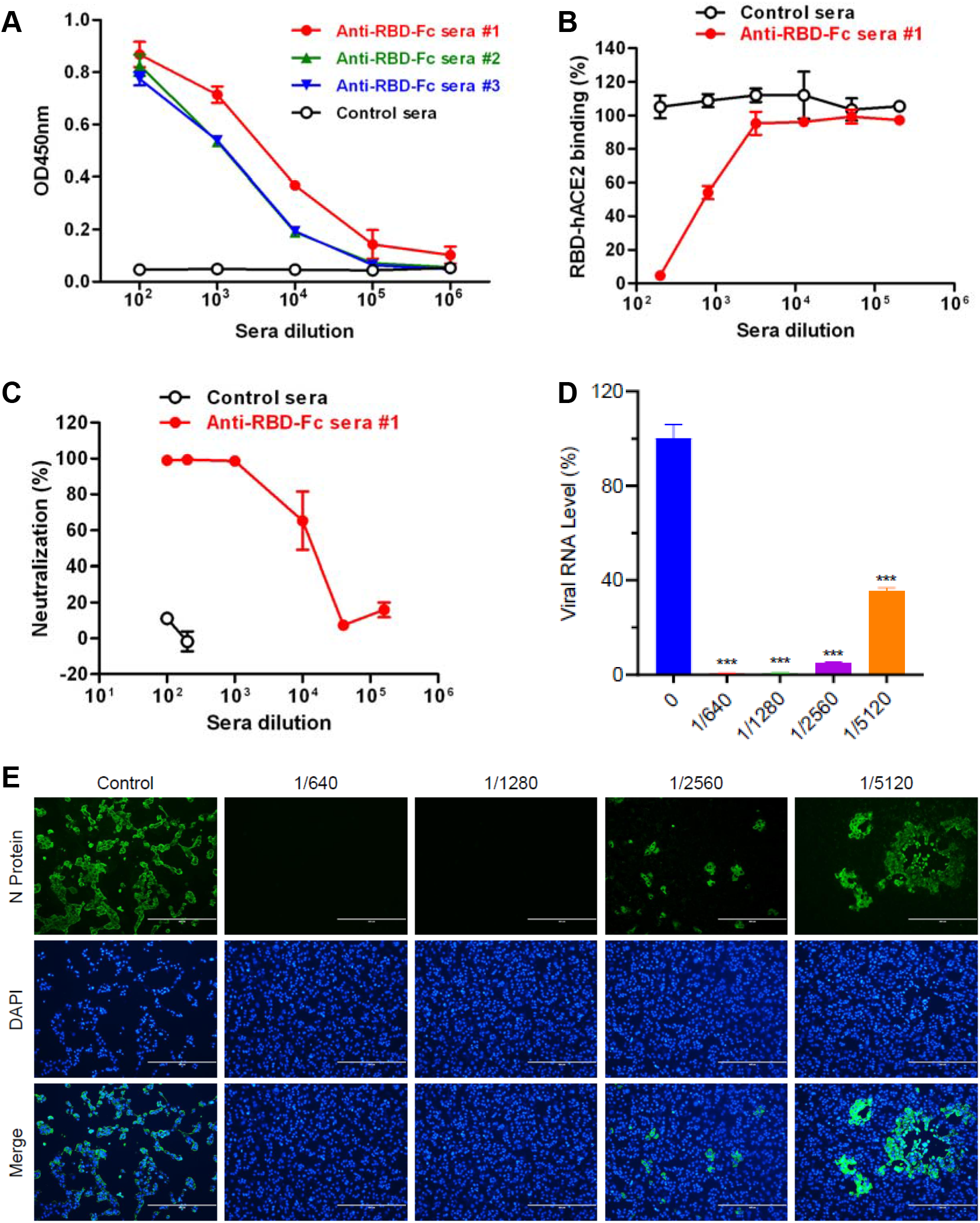
Immunization with recombinant RBD-Fc fusion protein potently elicited SARS-CoV-2 neutralizing antibodies in mice. (**A**) RBD-binding activities of the sera from the three RBD-Fc-immunized mice and the control (naïve) mouse. The sera were serially diluted and then analyzed by ELISA with recombinant SARS2-RBD protein as the coating antigen. Data shown are means and SD of triplicate wells. (**B**) Inhibitory effect of the anti-RBD-Fc sera on the RBD/ACE2 interaction. The anti-RBD-Fc sera #1 and the control sera were serially diluted and then subjected to ACE2 competition ELISA. Data shown are means and SD of triplicate wells. (**C**) Neutralization potency of the antisera against SARS-CoV-2 pseudovirus infection. The antisera were serially diluted and then evaluated for neutralization of SARS-CoV-2 spike-pseudotyped retrovirus. Results from three independent experiments are shown. (**D**) Neutralization potency of the antisera against authentic SARS-CoV-2 infection. Serially diluted antisera were subjected to neutralization assay with SARS-CoV-2 strain nCoV-SH01 as the challenge virus. Data shown are means and SD of triplicate wells. Significant differences were calculated using student’s two-tail t test and shown as: ***, P < 0.001. (**E**) Neutralization of authentic SARS-CoV-2 infection revealed by immunofluorescent staining. Live SARS-CoV-2 virus was incubated with or without serially diluted anti-RBD-Fc sera for 1 hr at 37°C and then added to preseeded VeroE6 cells. After three days, the cells were fixed and then stained sequentially with the N protein-specific primary antibody and a corresponding secondary antibody. Prior to examination under a fluorescent microscope, the cells were briefly stained with DAPI. Representative images are shown.

### RBD alone potently elicited cross-reactive, ACE2-binding blockade antibodies towards SARS-CoV-2 and SARS-CoV

To verify that the RBD part within the RBD-Fc fusion protein is indeed responsible for the induction of neutralizing antibodies against SARS-CoV-2, we performed a second mouse immunization study with recombinant SARS-CoV-2 RBD fused with a small 6xHis tag as the vaccine antigen. A group of six BALB/c mice were immunized by i.p. injection with 5 μg of recombinant RBD protein formulated with Alum adjuvant at days 1, 10, and 25 (Fig. 2A). Another group of mice were injected with an irrelevant protein (HBc) plus Alum adjuvant, serving as the control. Serum samples were collected from individual mice at days 20 and 40, and analyzed for RBD-specific antibody by ELISA using SARS2-RBD as the capture antigen. As shown in Fig. 2B, neither the day-20 nor the day-40 sera in the control (HBc) group exhibited any significant binding activity; in contrast, SARS2-RBD-binding activity was readily detectable at day 20 in the sera from RBD-immunized mice and a significant increase in SARS2-RBD-binding was observed for the day-40 anti-RBD sera. Equal amount of individual antisera in the same groups were pooled for antibody titer measurement and subsequent analyses. The day-20 and day-40 pooled anti-RBD sera dose-dependently reacted with SARS2-RBD in ELISAs (Fig. 2C) and their binding antibody titers were determined to be 1.6×10^5^ and 3.2×10^6^, respectively. The SARS2-RBD-binding activity of the anti-RBD sera collected at day 60 (when the mice were euthanized) was comparable to that of the day-40 anti-RBD sera (Supplemental Fig. 2).

**Figure 2.**
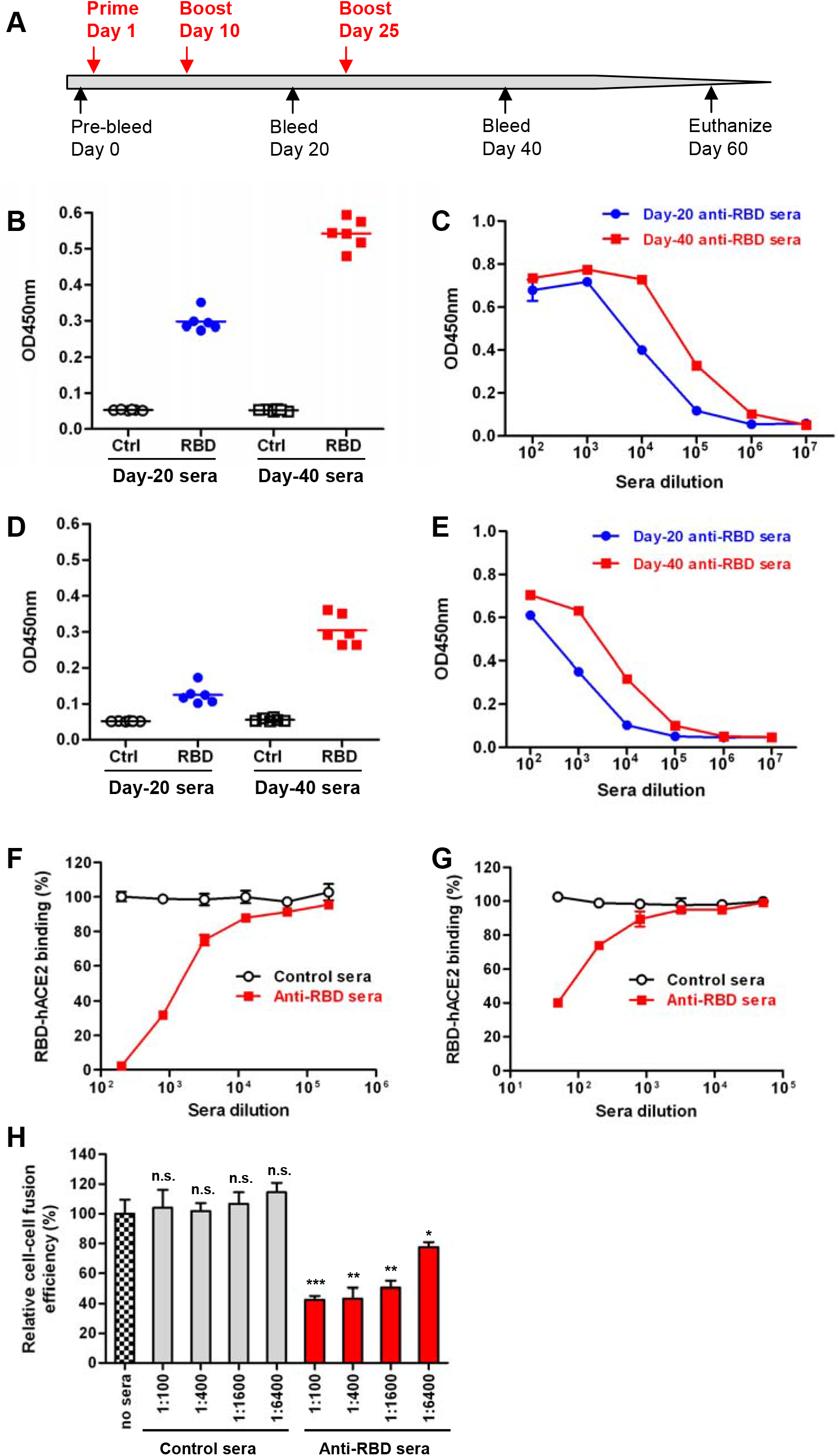
Mouse immunization with recombinant RBD protein and Characterization of the anti-RBD mouse sera. (**A**) Mouse immunization and sampling schedule. Two groups of six Balb/c mice received three doses of the RBD vaccine or the control antigen (HBc) on days 1, 10, and 25, respectively. The immunized mice were bled on days 20 and 40 and euthanized on day 60 (**B**) SARS2-RBD-binding activities of the individual antisera. The day-20 and day-40 serum samples were diluted 1:10,000 and then analyzed by ELISA with SARS2-RBD protein as the coating antigen. Each symbol represents a mouse and the line indicates geometric mean value of the group. (**C**) SARS2-RBD-binding activities of the pooled anti-RBD antisera. The indicated antisera were serially diluted and subjected to ELISA with SARS2-RBD protein as the coating antigen. Data shown are means and SD of triplicate wells. (**D**) Cross-reactivity of the individual antisera with recombinant SARS-RBD protein. The day-20 and day-40 serum samples were diluted 1:10,000 and then analyzed by ELISA with SARS-RBD as the coating antigen. Each symbol represents a mouse and the line indicates geometric mean value of the group. (**E**) SARS-RBD-binding activities of the pooled anti-RBD antisera. The same antisera as in (C) were analyzed by ELISA with SARS-RBD as the coating antigen. Data shown are means and SD of triplicate wells. (**F**) Blockade of ACE2 binding to SARS2-RBD by the anti-RBD sera. The pooled antisera were serially diluted and then subjected to ACE2 competition ELISA with SARS2-RBD as the capture antigen. Data shown are means and SD of triplicate wells. (**G**) Blockade of ACE2 binding to SARS-RBD by the anti-RBD sera. The pooled antisera were serially diluted and then subjected to ACE2 competition ELISA with SARS-RBD as the capture antigen. Data shown are means and SD of triplicate wells. (**H**) Inhibition of SARS2-S-mediated cell-cell fusion by the anti-RBD sera. HEK 293T cells transiently expressing SARS2-S:EGFP fusion protein were incubated with serially diluted antisera for 1 hr at 37°C and then mixed with HEK 293T cells transiently expressing hACE2:mCherry, followed by co-culture for 24 hours. The cells were analyzed by flow cytometry and the numbers of single- and dual-fluorescence cells were determined. For a given sample, its cell-cell fusion efficiency (the ratio of the dual-fluorescence cells to the EGFP-only cells) was normalized against that of the sample without antisera treatment. Data shown are means and SD of triplicate wells. Significant differences between the mock-treated (no sera) group and each of the antisera treatment groups were indicated: n.s., P>0.05; *, P < 0.05; **, P < 0.01; ***, P < 0.001.

SARS2-RBD shares high homology with the RBD of SARS-CoV (hereafter referred as SARS-RBD) in amino acid sequence, especially the regions outside the receptor-binding motif. This prompted us to evaluate the cross-reactivity of our SARS2-RBD-immunized sera towards SARS-RBD. As shown in Fig. 2D and 2E, both individual and pooled sera from the SARS2-RBD-immunized mice showed dose-dependent binding activity with SARS-RBD. The SARS-RBD-binding titers of the pooled day-20 and day-40 anti-RBD sera were determined to be 4×10^3^ and 1.6× 10^5^, respectively.

The pooled day-40 antisera were assessed for their ability to block the interaction between RBDs and ACE2, the entry receptor for both SARS-CoV-2 and SARS-CoV. Results from ACE2-binding blockade ELISA showed that the day-40 anti-RBD sera dose-dependently inhibited hACE2-Fc binding to SARS2-RBD whereas the control sera had no inhibitory effect on the SARS2-RBD/hACE2-Fc interaction regardless of the sera concentration used (Fig. 2F). The anti-RBD sera also exhibited blockade effect on the SARS-RBD/hACE2-Fc interaction, albeit with a lower efficiency (Fig. 2G).

### Anti-RBD sera inhibited SARS-CoV-2 spike-mediated cell-cell fusion

The spike protein of SARS-CoV-2 has been shown to bind cell surface ACE2 and mediate cell-cell fusion, leading to syncytia formation (Wang et al., 2020a). A cell-cell fusion assay was developed to determine whether the anti-RBD sera could prevent S-mediated syncytia formation. Co-culture of the 293T cells transiently expressing S:EGFP fusion protein and the ones expressing human ACE2 protein fused with mCherry (hACE2:mCherry) for 24 hours led to the detection of dual-fluorescent cells, indicating the occurrence of cell-cell fusion (Supplemental Fig. 3). The cells solely emitting green or red fluorescence and the dual-fluorescent cells were quantified by flow cytometry. It was found that addition of the day-40 anti-RBD antisera to the co-cultures significantly inhibited cell-cell fusion in an antisera dose-dependent manner whereas the control sera did not exhibit significant inhibitory effect regardless of the antisera dose used (Fig. 2H and Supplemental Fig. 4). This data indicate that the anti-RBD sera are able to inhibit SARS2-S-mediated cell-cell fusion.

### Anti-RBD sera neutralized both SARS-CoV-2 and SARS-CoV pseudoviruses

The neutralization capacity of the mouse antisera was first evaluated using SARS-CoV-2 S-pseudotyped retrovirus (SARS2-PV). The day-40 anti-RBD sera potently inhibited infection of hACE2-overexpressing VeroE6 cells (VeroE6-hACE2) with SARS2-PV and the calculated NT50 was 12764 (Fig. 3A), whereas the control sera did not show significant inhibition even at the lowest dilution tested (1:100). The same anti-RBD sera also inhibited infection of VeroE6-hACE2 cells with the SARS-PV (retrovirus pseudotyped with SARS-CoV S) with NT50 being 834.8 (Fig. 3B).

**Figure 3.**
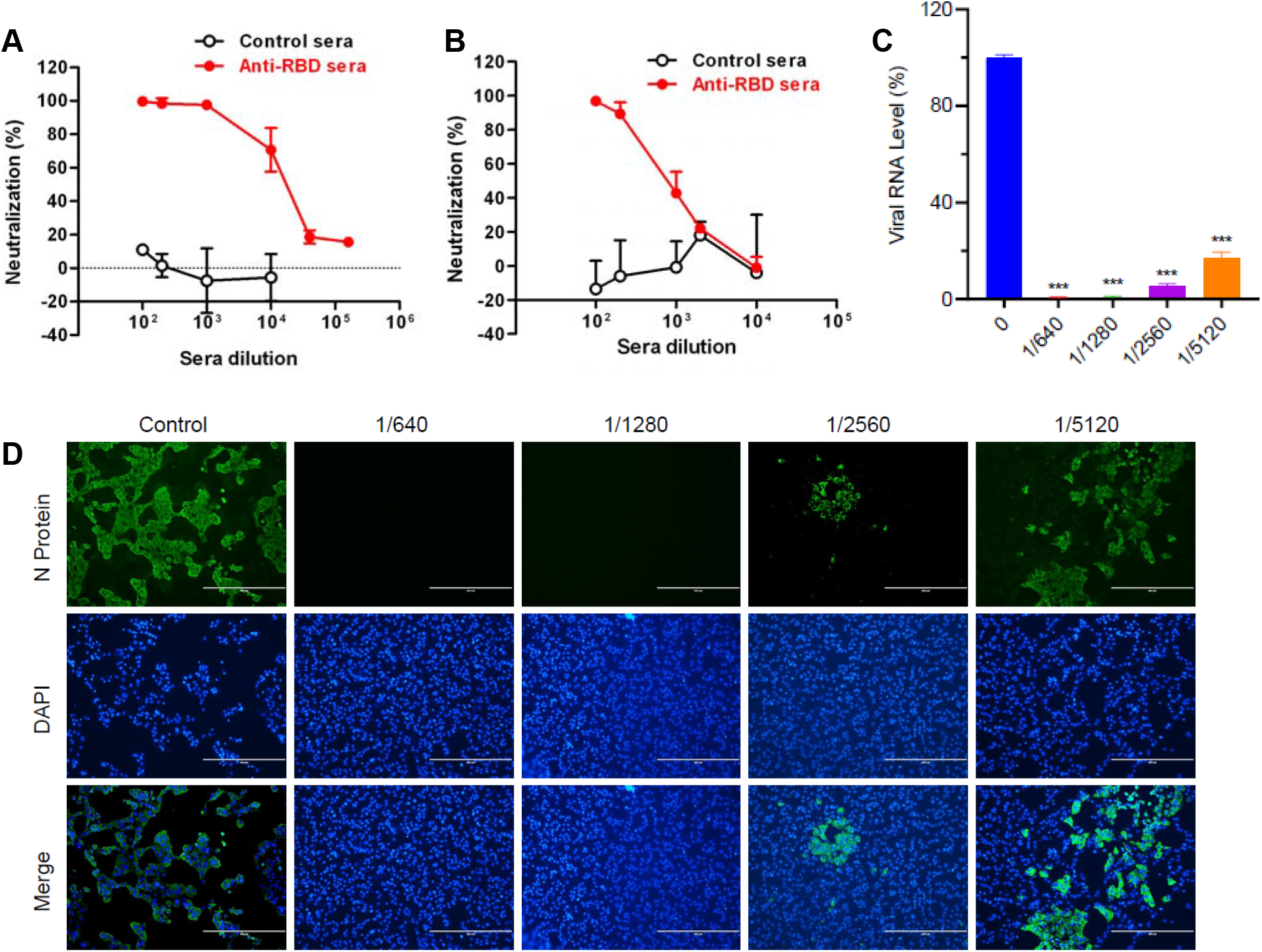
Neutralization potency and breadth of the anti-RBD sera. (**A**) Anti-RBD sera neutralized SARS2-PV infection in vitro. The day-40 pooled sera were serially diluted and tested for neutralization of retrovirus pseudotyped with SARS-CoV-2 S protein. Data (means±SD) from three independent experiments are shown. (**B**) Anti-RBD sera cross-neutralized SARS-PV infection in vitro. The day-40 pooled sera were serially diluted and tested for neutralization of retrovirus pseudotyped with SARS-CoV S protein. Data (means±SD) from three independent experiments are shown. (**C**) Neutralization efficiency of the anti-RBD sera against authentic SARS-CoV-2 infection. Serially diluted anti-RBD sera were mixed with 200 PFU of live SARS-CoV-2 and then incubated for 1 hr at 37°C. The antisera/virus mixtures were added to pre-seeded VeroE6 cells, followed by incubation for three days. The cells were then analyzed for viral RNA copy number by qPCR analysis. Data are expressed as percentage of the viral RNA copy number of the treatment groups in relation to that of the virus-only control. Means ± SD of triplicate wells are shown. Significant differences between treatment groups and the virus-only control were calculated using student’s two-tail t test and shown as: ***, P < 0.001. (**D**) Neutralization of authentic SARS-CoV-2 infection revealed by immunofluorescent staining. Representative images are shown.

### Anti-RBD sera neutralized authentic SARS-CoV-2 infection

The antisera were next tested for neutralization of authentic SARS-CoV-2 infection. The anti-RBD sera were found to potently inhibite infection of VeroE6 cells with authentic SARS2-CoV-2 based on CPE observation (Supplemental Fig. 5), whereas the control sera did not prevent CPE development even when a sera dilution of 1:20 (the lowest dilution tested) was used. Both qRT-PCR and IFA assays revealed that the anti-RBD sera diluted 1:1280 nearly completely block the viral infection and even the 1:5120 diluted anti-RBD sera could inhibit viral infection by 83% (Fig. 3C-D). These results demonstrate that the anti-RBD sera possessed very strong neutralization capacity against SARS-CoV-2.

### Anti-RBD mouse sera did not enhance SARS-CoV-2 infection in Fc receptor-expressing cells

For some viruses such as dengue virus, antibodies targeting the envelope protein may increase viral infection of FcR-expressing cells – a phenomenon called antibody-dependent enhancement (ADE) (Huisman et al., 2009; Thomas et al., 2006). Several FcR-bearing cell lines were used as the target cell in infection/neutralization assays to evaluate whether our antisera could mediate ADE in vitro, including mouse A20 cells expressing FcγRII (Antoniou and Watts, 2002), human THP-1 cells expressing both FcγRI and FcγRII (Chan et al., 2011), and K562 cells expressing human FcγRII (Block et al., 2010). Both THP-1 and K562 cells have been shown to support mouse antibody-mediated enhancement of dengue virus infection in previous studies (Block et al., 2010; Sun et al., 2017; Zhang et al., 2019). We found that SARS2-PV entry into the three FcR-expressing cell lines was minimal (<0.02%) whereas the same amount of SARS2-PV yielded an infection rate of ~7% in VeroE6-hACE2 cells. Moreover, treatment with serially diluted (ranging from 1:10^2^ to 1:10^6^) control sera or anti-RBD sera did not significantly affect SARS2-PV entry of the three cell lines (Fig. 4A-C), indicating that the anti-RBD sera do not promote ADE of SARS2-PV. We selected K562 cells for performing ADE assay with authentic SARS-CoV-2 as the challenge virus. No significant increase in viral RNA level was observed for the antisera-treated samples as compared to the control (cells only infected with the virus) regardless of the antisera dilutions (Fig. 4D). Collectively, these results demonstrate that, in the assay system we tested, the anti-RBD antibodies do not promote ADE.

**Figure 4.**
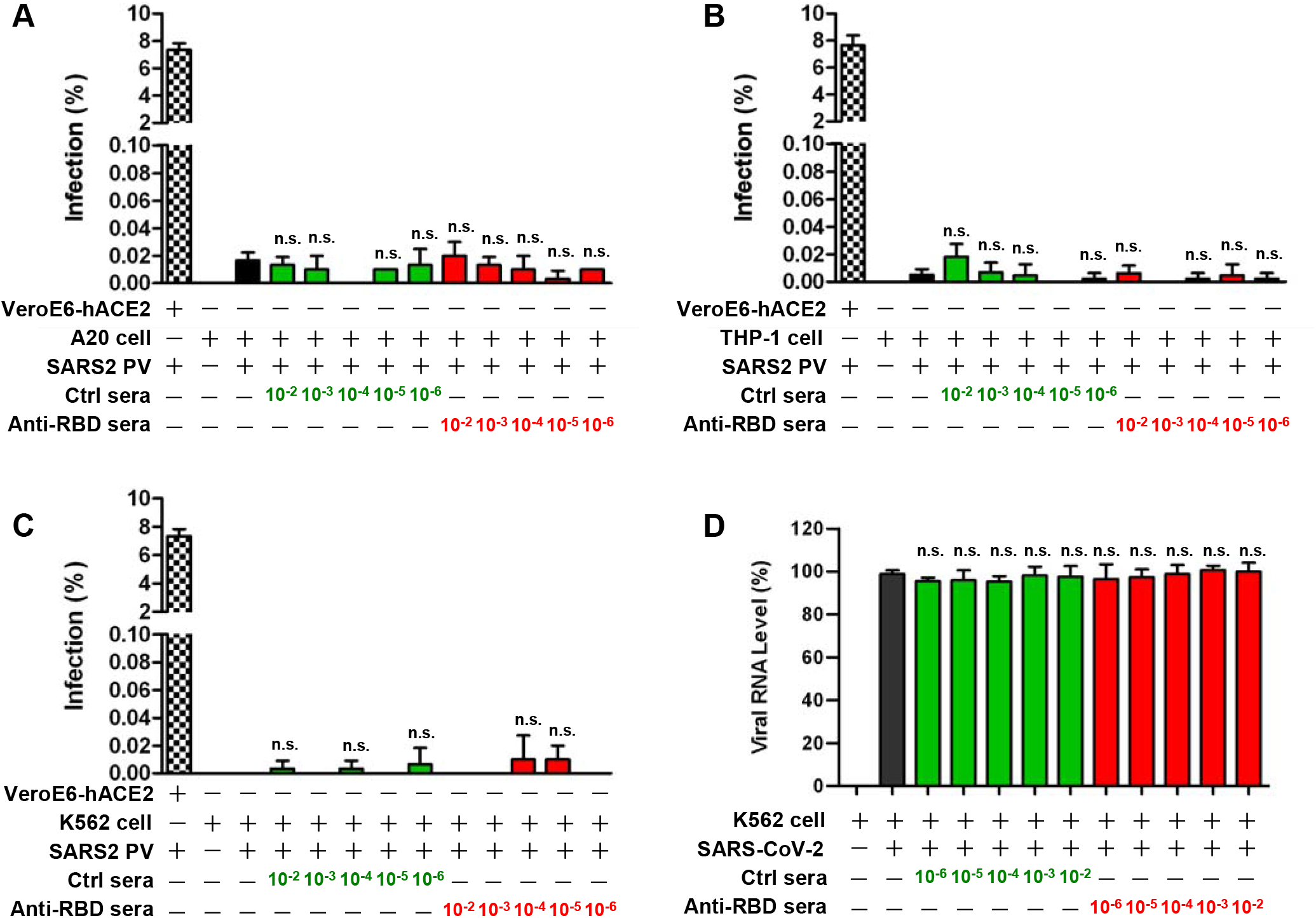
Assessment of the anti-RBD sera for potential ADE. (**A-C**) ADE assays with SARS2-PV as the inoculum. Serial dilutions of the anti-RBD or the control sera were incubated with SARS2-S pseudotyped retrovirus for 1 hour at 37 °C. The mixtures were added to (**A**) A20, (**B**) THP-1, or (**C**) K562 cell suspensions, followed by incubation at 37°C for three days. Infected cells were subjected to flow cytometry analysis. Data are expressed as percentage of the GFP-expressing cells in relation to the total cells counted. Means ± SD of triplicate wells are shown. (**D**) ADE assay with live SARS-CoV-2 virus as the inoculum. Serial dilutions of the anti-RBD or the control sera were mixed with the live SARS-CoV-2 virus and incubated for 1 hour at 37 °C. The mixtures were added to K562 cell suspensions, followed by incubation at 37 °C for three days. Infected cell cultures were subjected to RNA extraction and qPCR analysis. Data are expressed as percentage of the viral RNA copy number of the treatment groups in relation to that of the virus-only control. Means ± SD of triplicate wells are shown. Significant differences between the virus only (without antisera treatment) group and each of the antisera treatment groups were indicated: n.s., P>0.05.

## DISCUSSION

The present study was aimed to investigate the possibility of developing a RBD-based subunit vaccine for SARS-CoV-2. It was found that RBD fusion proteins with either Fc or 6 × His tag elicited high-titer neutralizing antibody responses, indicating that RBD is the antigen component responsible for the induction of neutralizing antibodies. In particular, the anti-RBD sera potently neutralized SARS2-PV entry with an NT50 value of 12764 (Fig. 3A) and its neutralization efficiency against authentic SARS-CoV-2 was >75% at the serum dilution of 1:5120 (Fig. 3C). According to a recent report (Gao et al., 2020), macaques immunized with a purified inactivated SARS-CoV-2 virus vaccine candidate (which induced NT50 titers up to 3,000 in mice) produced neutralizing serum antibody titers of around 50 to 61 and were protected from authentic SARS-CoV-2 challenge. Therefore, the seemingly higher neutralizing antibody titers of our anti-RBD mouse antisera compared to that of the mouse antisera against the inactivated whole-virus vaccine (although not directly compared) strongly suggest that the RBD subunit vaccine candidate will likely be protective as well.

The anti-RBD sera not only were able to efficiently neutralize SARS-CoV-2 infections but also could inhibit SARS2-S-mediated cell-cell fusion (Fig. 2H and Supplementary Fig. S3 and S4), demonstrating a two-layer protective potential of the antisera. Host receptor recognition and binding is required for coronavirus entry. Previous work showed that, for SARS-CoV and MERS-CoV, RBD binding to the receptor triggers the spike to undergo conformational changes, allowing subsequent protease cleavage, virus internalization and membrane fusion (Millet and Whittaker, 2018; Pallesen et al., 2017). In this study, the anti-RBD sera were found to dose-dependently reduce the RBD binding to ACE2 in ELISAs (Fig. 1B and 2F). These data suggest blocking the interaction between RBD/S and the host receptor ACE2 is the main mechanism underlying the observed neutralization and cell-cell fusion inhibition by the anti-RBD sera.

Interestingly, we found that the mouse antisera raised against SARS2-RBD cross-reacted with SARS-RBD, inhibited SARS-RBD binding to ACE2, and neutralized SARS-PV entry with an NT50 of 834.8. SARS2-RBD can be divided into a core subdomain and a receptor-binding motif (RBM) which directly engages ACE2 (Lan et al., 2020; Shang et al., 2020; Yan et al., 2020). The core subdomain is highly conserved while the RBM varies significantly (~47% homology in amino acid sequence) between SARS-CoV and SARS-CoV-2. Therefore, we reason that the observed cross binding and neutralization capacity towards SARS-CoV is contributed by antibodies targeting the conserved core subdomain of SARS2-RBD. That is to say, the core subdomain contains SARS-CoV-2/SARS-CoV cross-neutralization antibody epitopes. This information will be useful for future design and development of pan-SARS-CoV vaccines.

A major concern in developing coronavirus vaccines is the risk of vaccine-induced ADE (Hotez et al., 2020; Huisman et al., 2009). ADE phenomenon has been observed for feline coronavirus (Huisman et al., 1998; Huisman et al., 2009; Weiss and Scott, 1981) and for SARS-CoV (Bolles et al., 2011; Luo et al., 2018; Tseng et al., 2012; Wang et al., 2016). Some cell culture studies suggested that anti-S antibodies mediated ADE in FcR-expressing cells likely through enhancing FcR-mediated internalization/entry of antibody-bound virions (Corapi et al., 1992; Jaume et al., 2011; Liu et al., 2019; Olsen et al., 1992; Wan et al., 2020; Wang et al., 2014; Yip et al., 2014). In the present study, we showed that the anti-RBD sera did not enhance SARS2-PV entry of the three FcR-expressing cell lines (A20, THP-1, and K562) regardless of the antisera concentration. Moreover, treatment with the anti-RBD sera did not increase authentic SARS-CoV-2 infection of K562 cell, despite the same cell line has been shown to support anti-DENV-E mouse antibody-triggered ADE of Dengue virus (DENV) infection in previous studies (Block et al., 2010; Sun et al., 2017; Zhang et al., 2019). These data clearly show that anti-RBD antibodies do not promote ADE, at least not in the assay system we used. It remains to be determined whether antibodies targeting other regions of the S protein (which presumably will be non-neutralizing or poorly neutralizing) could mediate ADE of SARS-CoV-2. Nonetheless, the extraordinary neutralization potency and lack of ADE effect observed for the anti-RBD sera indicate that SARS2-RBD is an elite antigen target for developing subunit vaccines for SARS-CoV-2.

Collectively, our results show that recombinantly expressed SARS2-RBD proteins potently elicits cross-neutralizing antibodies against both SARS-CoV-2 and SARS-CoV without induction of ADE antibodies, providing important information for further development of RBD-based SARS-CoV-2 or pan-SARS-CoV subunit vaccines.

## MATERIALS AND METHODS

### Cells and viruses

VeroE6 cells were grown as described previously (Zhao et al., 2018). HEK 293T, A20, THP-1, and K562 cells were purchased from the Cell Bank of Chinese Academy of Sciences (www.cellbank.org.cn). A clinical isolate of SARS-CoV-2, nCoV-SH01 (GenBank: MT121215.1) (Rong et al., 2020), was propagated in VeroE6 cells and viral titer was determined as plaque forming units (PFU) per milliliter (mL) by CPE quantification. Live virus infection experiments were performed in the biosafety level-3 (BSL-3) laboratory of Fudan University.

### Recombinant proteins

For mouse immunization, recombinant SARS-CoV-2 RBD fusion protein with the Fc region of mouse IgG1 at the C-terminus (RBD-Fc) was purchased from Sino Biological (Beijing, China), recombinant SARS-CoV-2 RBD with a C-terminal His-tag was purchased from Kactus Biosystems (Shanghai, China), and recombinant hepatitis B core antigen (HBc) was produced in house in *E.coli* as described previously (Ye et al., 2014). For biochemical and immunological assays, several mammalian cell-produced recombinant proteins were generated in house, including SARS-CoV-2 RBD (amino acids 320 to 550) fused with an N-terminal Strep-tag and a C-terminal His-tag, SARS-CoV RBD (amino acids 306-520) fused with a C-terminal His-tag, and human ACE2 ecotodomain fused with human IgG1 Fc at the C-terminus (hACE2-Fc). Biotinylated hACE2-Fc was prepared using EZ-Link™ Sulfo-NHS-LC-LC-Biotin kit (Thermo Fisher Scientific).

### Mouse immunization

All the animal experiments in this study were approved by the Institutional Animal Care and Use Committee at the Institut Pasteur of Shanghai. Animals were cared for in accordance with institutional guidelines.

In the first immunization experiment, three BALB/c mice were each injected intraperitoneally (i.p.) with 100 μg of RBD-Fc fusion protein formulated with 500 μg of aluminum hydroxide (Alhydrogel, Invivogen, USA) and 25 μg of CpG (Sangon, China) at day 0. The mice were boosted subcutaneously (s.c.) at day 8 with 50 μg of RBD-Fc plus Freund’s Adjuvant Complete (Sigma, USA) and at day 13 with 50 μg of RBD-Fc plus Titermax adjuvant (Sigma). Blood were collected from individual mice one week after the last immunization (day 20) and sera were stored at −80°C until use.

In the second immunization experiment, recombinant RBD protein containing a C-terminal 6xHis tag was formulated with the Alhydrogel adjuvant (Invivogen) and each vaccine dose contained 5 μg of RBD and 500 μg of aluminum hydroxide; in addition, a negative antigen control was prepared by mixing 5 μg of recombinant HBc protein with 500 μg of aluminum hydroxide. Two groups of six BALB/c mice were injected intraperitoneally (i.p.) with the RBD vaccine and the control antigen, respectively, at days 0, 10, and 25. Blood samples were collected from individual mice at days 20, 40 and 60 for antibody measurement.

### Serum antibody measurement

For antibody measurement, wells of 96-well microtiter plates were coated with the indicated amounts of the SARS2-RBD or SARS-RBD recombinant protein for 2 hrs at 37°C or overnight at 4°C. Then the wells were blocked with PBST containing 5% non-fat dry milk for 1 hr at 37°C, incubated with 50μl serially diluted mouse antisera for 2 hrs at 37°C and then with 50 μl of horseradish peroxidase (HRP)-conjugated goat anti-mouse IgG antibody for 1 hr 37°C. After color development, the absorbance at 450 nm was measured in a 96-well plate reader. For a given serum sample, its endpoint titer was reported as the reciprocal of the highest serum dilution that had an absorbance ≥0.1 OD unit above the blank.

### ACE2 competition ELISA

Wells of 96-well microtiter plates were coated with 25 ng/well of the SARS2-RBD or SARS-RBD recombinant protein overnight at 4°C, followed by blocking with PBST containing 5% non-fat dry milk for 1 hr at 37°C. Serially diluted mouse antisera were mixed with 20 ng of biotinylated hACE2-Fc in a final volume of 50μl and the mixtures were added to the wells, followed by incubation for 2 hrs at 37°C. Then, 50 μl of horseradish peroxidase (HRP)-conjugated streptavidin (Life Technologies, USA) was added to wells, followed by incubation for 1 hr at 37°C. After washing, TMB substrate (Life Technologies) was added into wells for color development. The plates were read for absorbance at 450 nm in a 96-well plate reader.

### Cell-cell fusion inhibition assay

HEK 293T cells were separately transfected with a plasmid encoding the SARS-CoV-2 S:EGFP fusion protein (pcDNA-S:EGFP) or with a plasmid encoding the hACE2:mCherry fusion protein (pcDNA-hACE2:mCherry). One day later, equal amount of the pcDNA-S:EGFP-transfected and pcDNA-hACE2:mCherry-transfected cells were mixed and then cultured for 24 hrs. Unmixed cells were set aside as controls. To determine the antisera’s blockade effects, pcDNA-S:EGFP-transfected cells were treated with serially diluted antisera for 1 hr at 37°C before mixing with pcDNA-hACE2:mCherry-transfected cells. After co-culture for 24 hrs, the cells were subjected to fluorescence microscopy or flow cytometry. The cells emitting green or red fluorescence only or both were quantified by flow cytometry. For a given sample, its cell-cell fusion efficiency was calculated and normalized against that of the sample without antisera treatment as follows: relative cell-cell fusion efficiency (%) = (the ratio of the dual-fluorescence cells to the EGFP-only cells of the given sample) / (the ratio of the dual-fluorescence cells to the EGFP-only cells of the sample without antisera treatment)×100.

### Pseudovirus neutralization assay

To produce pseudoviruses, HEK293T cells were transfected using PEI with a plasmid encoding murine leukemia virus (MLV) gag/pol, a retroviral vector encoding EGFP, and an envelope plasmid expressing full-length S protein of SARS-CoV-2 or SARS-CoV (AY569693). Six hours later, the cells were washed and incubated in fresh medium. At 48 hours post-transfection, pseudovirus-containing culture supernatants were harvested. For neutralization assay, 100 μl of the pseudovirus was pre-mixed with 50 μl of serum samples diluted in DMEM and incubated at 37°C for 1 hr. The mixture was then onto VeroE6 cells overexpressing hACE2 (denoted as VeroE6-hACE2) preseeded in 48-well plates. Eight hours later, the virus/sera-containing media were removed and exchanged with fresh media containing 10% FBS. At 72 hours post-infection, the cells were analyzed by flow cytometry. The infectivity of pseudotyped particles incubated with antibodies was compared with the infectivity observed using pseudotyped particles incubated with DMEM medium containing 2% fetal calf serum (FBS) and standardized to 100%.

### Live virus neutralization assay

All serum samples were heat-inactivated at 56°C for 30 minutes prior to live virus neutralization assay. SARS-CoV-2 virus (200 PFU in a volume of 50μl) was pre-incubated with the diluted serum sample for 1 hour at 37°C. The virus-serum mixture was then added onto VeroE6 cells (4×10^4^/well) in 96-well plate and cultured for 48 hours. At the end of the incubation, culture supernatants were collected for viral RNA analysis and cells were fixed for immunofluorescence analysis.

Viral RNA in culture supernatant was extracted using TRIzol reagent (Invitrogen, USA) following the manufacturer’s instructions. Quantitative real-time PCR (qRT-PCR) was performed in a 20-μL reaction containing SYBR Green (Tiangen, China) on an MXP3000 cycler (Stratagene, La Jolla, USA). PCR primers (Genewiz, Suzhou, China) targeting SARS-CoV-2 N gene (nt 608-706) were as the following: forward primer, 5’-GGGGAACTTCTCCTGCTAGAAT-3’; and reverse primer, 5’-CAGACATTTTGCTCTCAAGCTG-3’.

For immunofluorescence analysis, cells were fixed in 4% paraformaldehyde, permeabilized by 0.2% Triton X-100 (Thermo Fisher Scientific, USA), and stained overnight at 4°C with an anti-N mouse polyclonal antibody generated in house. The samples were finally incubated with Alexa Fluor 488-labeled donkey anti-mouse IgG secondary antibody (1:1000, Thermo Fisher Scientific) at 37°C for 1 hour. The nuclei were stained with DAPI (Thermo Fisher Scientific). Images were captured under a fluorescence microscope (Thermo Fisher Scientific).

### ADE assay

FcR-expressing cell lines, including A20, THP-1, and K562, were used to perform ADE assays. Briefly, the antisera were serially diluted, mixed with either SARS2-PV or authentic SARS-CoV-2 (6,000 PFU), and incubated at 37°C for 1 hr. Then, the mixtures were added to the target cells. The following infection and culturing steps were carried out as described above in the pseudovirus neutralization and live virus neutralization assays. Mock-infected cells and cells only infected with SARS2-PV or authentic SARS-CoV-2 were set as the negative and positive controls, respectively. Infection rates of the samples were determined as described above.

### Statistics analysis

All statistical analyses were performed using GraphPad Prism software v5.0. Kaplan–Meier survival curves were compared using log-rank test. Statistical significance between treatments was analyzed using Student’s 2-tailed *t*-test and indicated as follows: ns, not significant (P ≥ 0.05); *, 0.01 ≤ P < 0.05; **, P < 0.01; ***, P < 0.001.

## ACKNOWLEDGEMENTS

We thank Dr. Xiaozhen Liang for providing A20 and K562 cell lines, Dr. Guangxun Meng for THP-1 cell line, Drs. Gary Wong and Jiaming Lan for codon-optimized S gene, and Dr. Haikun Wang for assistance in flow cytometry analysis. This study was supported by grants from the Chinese Academy of Sciences (XDB29040300) and from the Chinese Ministry of Science and Technology (2020YFC0845900). The BSL-3 lab of Fudan University was supported by Shanghai Science and Technology Committee and Project of Novel Coronavirus Research from Fudan University.

---

**Supplemental Figure 1.**
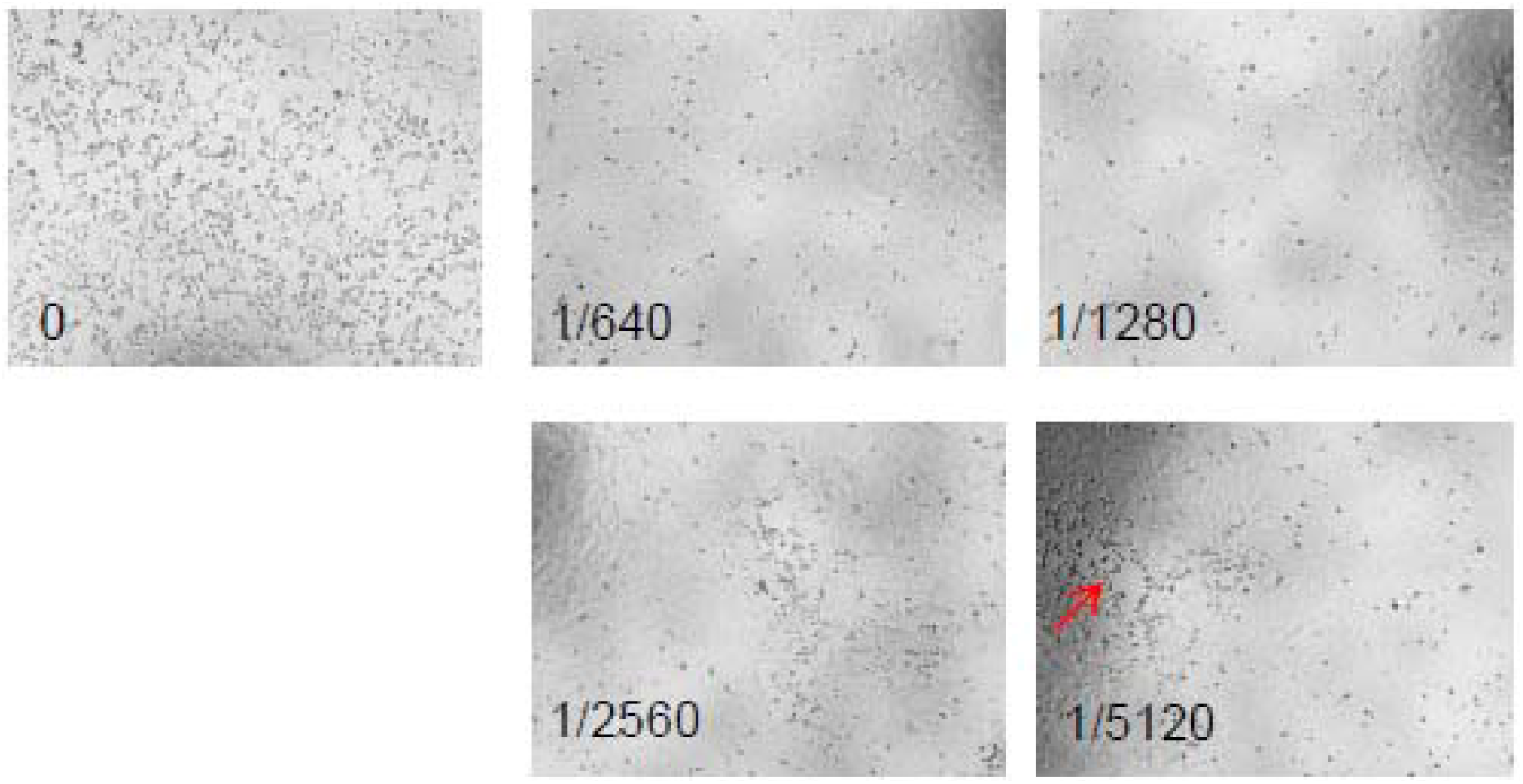
Treatment with the anti-RBD-Fc sera #1 inhibited SARS-CoV-2 infection-triggered CPE. VeroE6 cells were inoculated with mixtures of authentic SARS-CoV-2 and serially diluted anti-RBD-Fc sera #1. The cells were checked daily for CPE. Data presented are images taken at 48 hours post-infection. The test concentrations of the anti-RBD-Fc sera #1 are indicated.

**Supplemental Figure 2.**
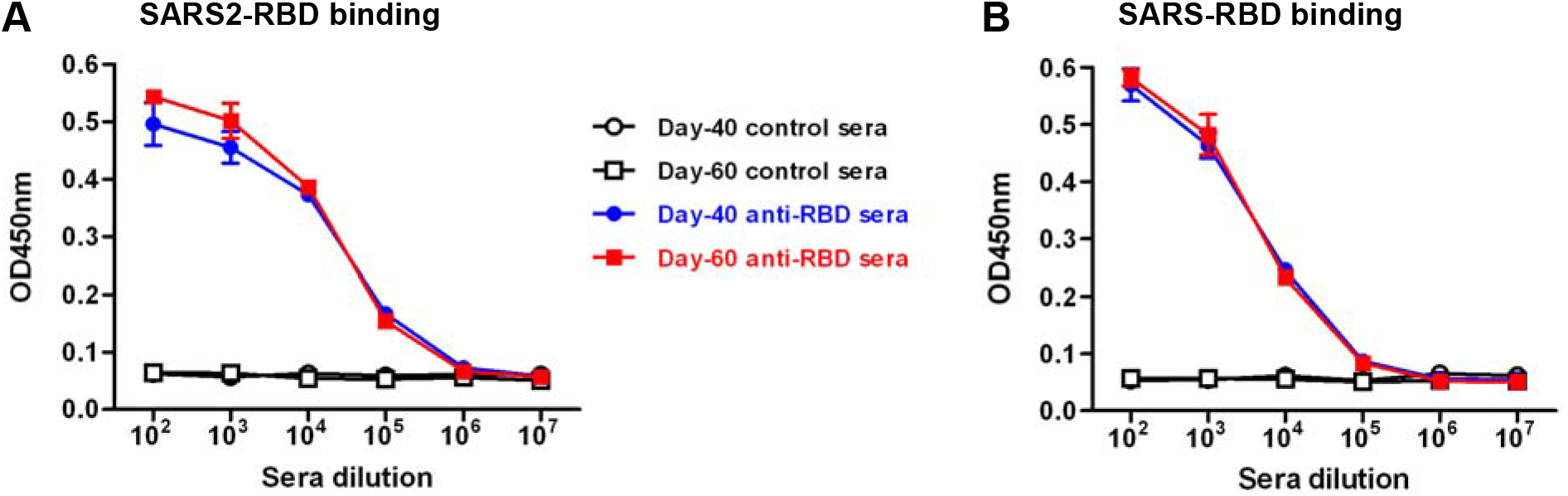
Comparison of binding activities of the day-40 and day-60 anti-RBD sera pools. The indicated antisera were serially diluted and analyzed by ELISA with (**A**) SARS2-RBD or (**B**) SARS-RBD proteins as the coating antigen. Data shown are mean OD450nm values and SD of triplicate wells.

**Supplemental Figure 3.**
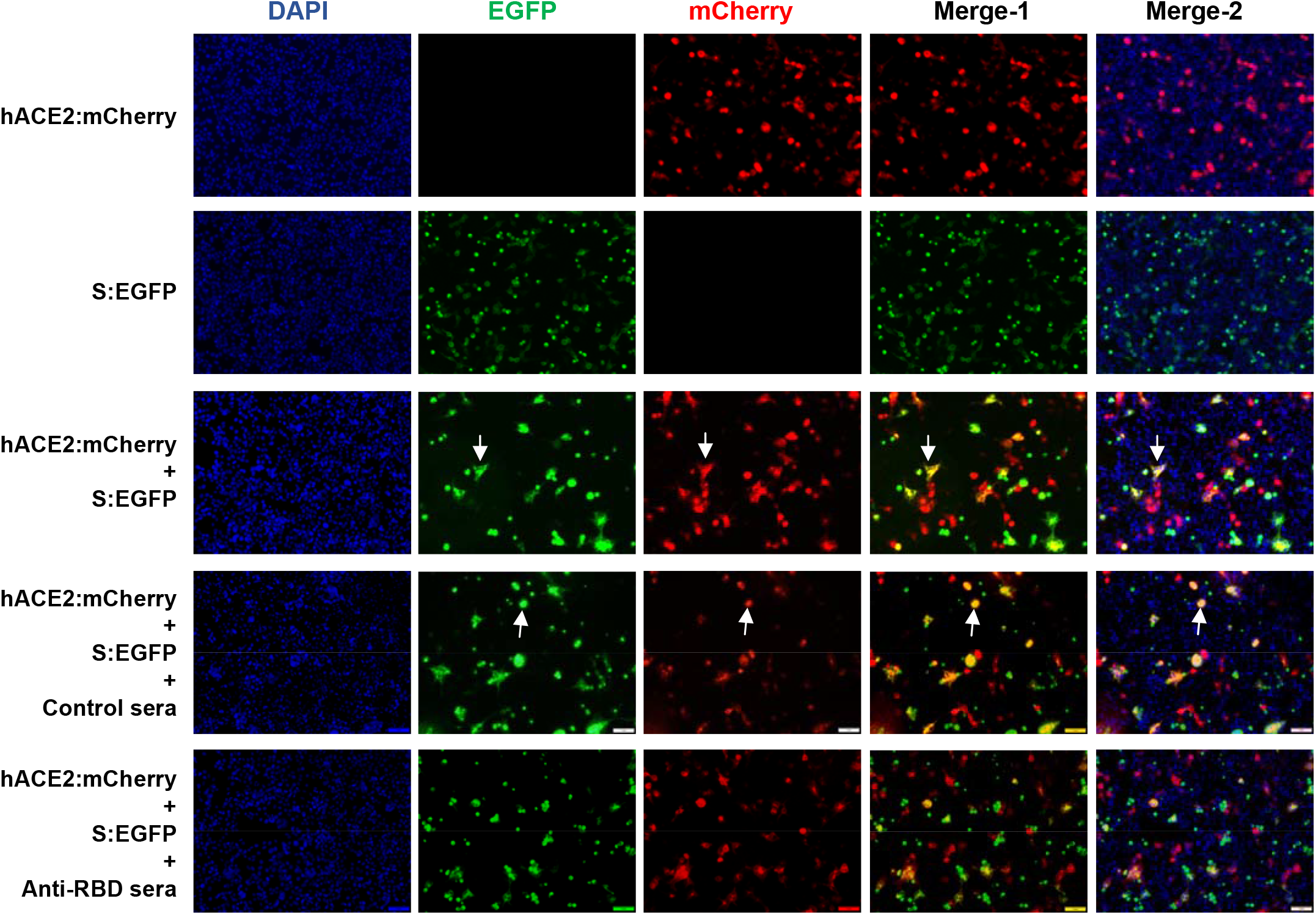
Cell-cell fusion assay. HEK 293T cells were separately transfected with a plasmid encoding the SARS2-S:EGFP fusion protein (pcDNA-S:EGFP) or with a plasmid encoding the hACE2:mCherry fusion protein (pcDNA-hACE2:mCherry). One day later, equal amount of the pcDNA-S:EGFP-transfected and pcDNA-hACE2:mCherry-transfected cells were mixed and then cultured for 24 hrs. Unmixed cells were set aside as controls. To determine the antisera’s blockade effects, pcDNA-S:EGFP-transfected cells were treated with serially diluted antisera for 1 hr at 37 °C before mixing with pcDNA-hACE2:mCherry-transfected cells. After co-culture for 24 hrs, the cells were fixed, stained with DAPI and examined under a fluorescent microscope. Representative images are shown. Blue signal, DAPI; green signal, S:EGFP; red signal, hACE2:mCherry; Merge 1, merge of the green and red channels; Merge 2, merge of the blue, green and red channels.

**Supplemental Figure 4.**
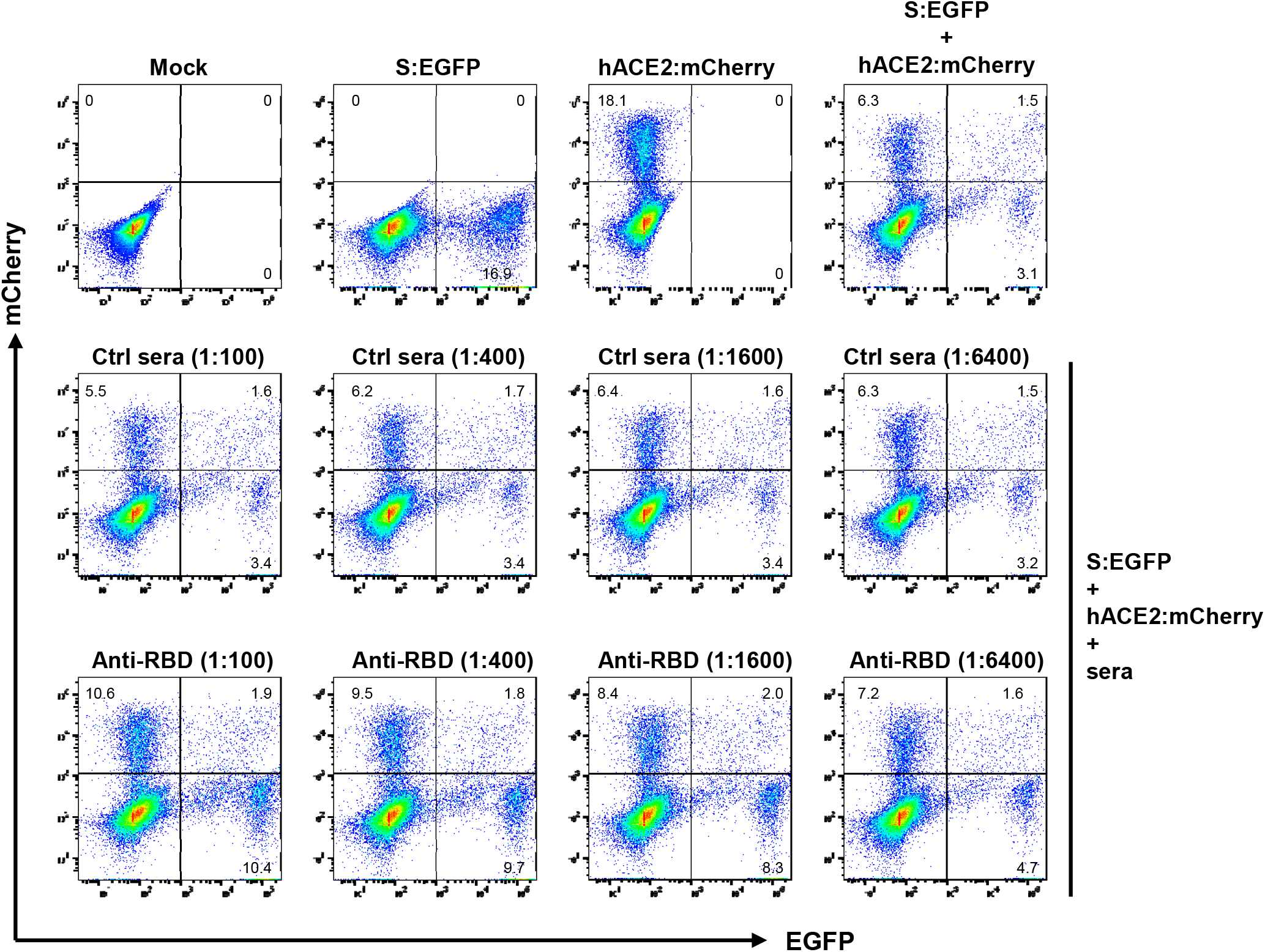
Inhibition of SARS2-S-mediated cell-cell fusion by anti-RBD sera. HEK 293T cells transiently expressing SARS2-S:EGFP fusion protein were incubated with the indicated dilutions of antisera for 1 hr at 37°C and then mixed with HEK 293T cells transiently expressing hACE2:mCherry, followed by co-culture for 24 hours. Cells without antisera treatment were set as the control. The cell samples were analyzed by flow cytometry. Representative flow cytometry graphs are shown. For a given sample, its cell-cell fusion efficiency was determined as the ratio of the dual-fluorescence cells to the EGFP-only cells

**Supplemental Figure 5.**
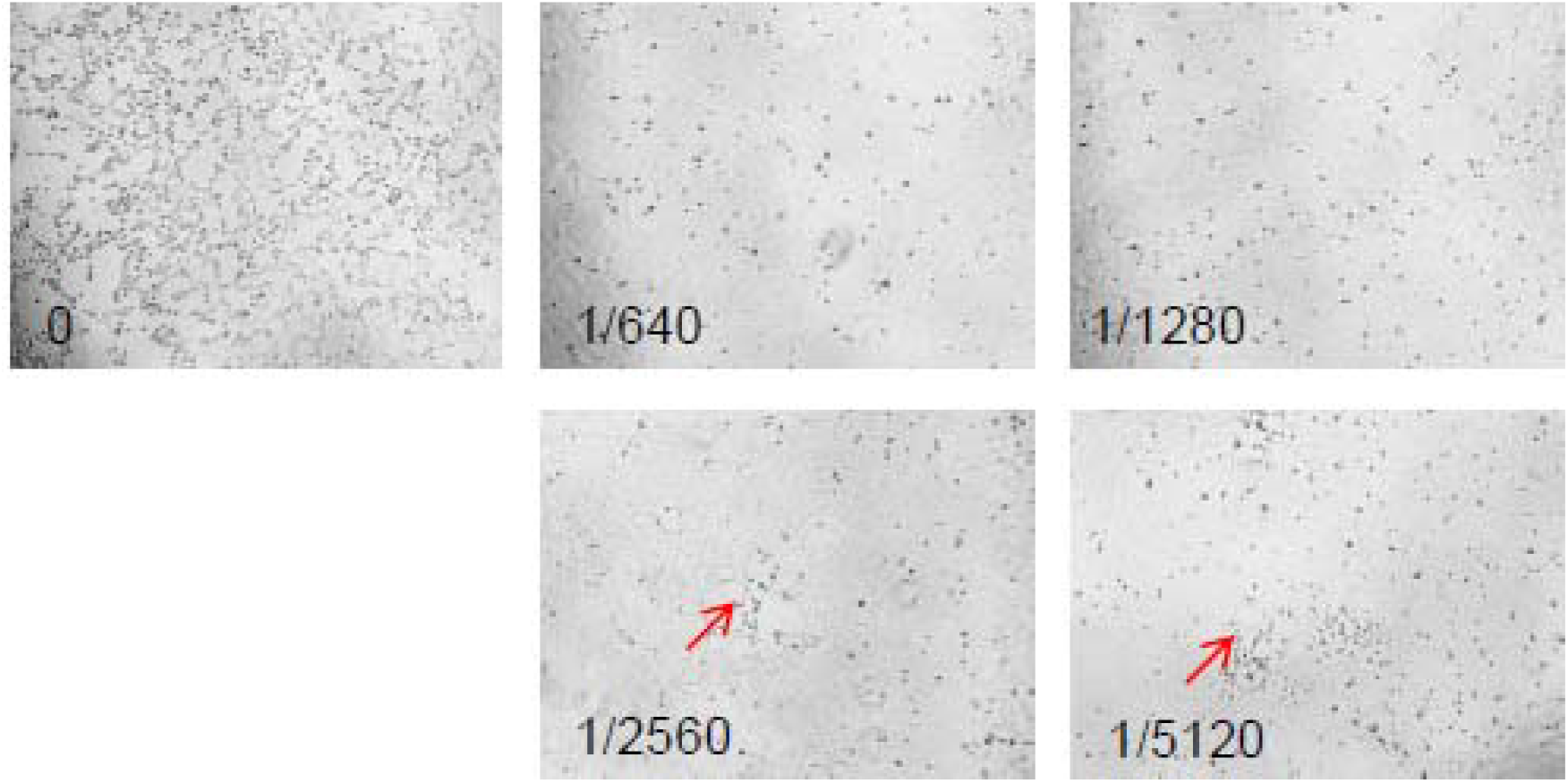
Treatment with the anti-RBD sera inhibited SARS-CoV-2 infection-triggered CPE. VeroE6 cells were inoculated with mixtures of the authentic SARS-CoV-2 virus and serially diluted anti-RBD sera. The cells were checked daily for CPE. Data presented are images taken at 48 hours post-infection. The test concentrations of the anti-RBD sera are indicated.

